# Time-dependent Canonical Correlation Analysis for Multilevel Time Series

**DOI:** 10.1101/650101

**Authors:** Xuefei Cao, Jun Ke, Björn Sandstede, Xi Luo

**Affiliations:** 182 George Street, Providence, RI; 121 S Main St, Level 7, Providence, RI; 1200 Pressler St, Houston, TX 77030

**Keywords:** Canonical correlation analysis, time series, temporal dynamics, fMRI

## Abstract

Canonical Correlation Analysis is a technique in multivariate data analysis for finding linear projections that maximize the correlation between two groups of variables. The correlations are typically defined without accounting for the serial correlations between observations, a typical setting for time series data. To understand the coupling dynamics and temporal variations between the two time-varying sources, we introduce the time-dependent canonical correlation analysis (TDCCA), a method for inferring time-dependent canonical vectors from multilevel time series data. A convex formulation of the problem is proposed, which leverages the singular value decomposition (SVD) characterization of all solutions of the CCA problem. We use simulated datasets to validate the proposed algorithm. Moreover, we propose a novel measure, canonical correlation variation as another way to assess the dynamic pattern of brain connections and we apply it to a real resting state fMRI dataset to study the aging effects on brain connectivity. Additionally, we explore our proposed method in a task-related fMRI to detect the temporal dynamics due to different motor tasks. We show that, compared to extant methods, the TDCCA-based approach not only detect temporal changes but also improves feature extraction. Together, this paper contributes broadly to new computational methodologies in understanding multilevel time series.

## 1. Introduction

Canonical Correlation Analysis (CCA) [1] is a powerful tool to analyze the relationship between two sets of variables. CCA can be regarded as an extension of ordinary correlation analysis, the difference being that CCA deals with multidimensional variables. It finds two linear transformations, one for each set of variables, that are optimal based on their correlations. It is an especially useful technique in data analysis as a dimensional reduction strategy that reduces the complexity of model space by calculating the combinations of variables that are maximally correlated. In an attempt to increase the flexibility for large dimensional date, several extensions of CCA have been proposed, including kernel Canonical Correlation Analysis (KCCA) [2, 3], Sparse Canonical Correlation Analysis (SCCA) [4, 5, 6, 7, 8]. Together, CCA-type methods have various applications including analysis of neuroimage, genomic data and information retrieval [9, 6, 10].

For multilevel data, CCA has been studied extensively by multi-view CCA [11] and tensor CCA [12]. However, canonical correlation analysis of multivariate longitudinal data with multiple observations has received considerably less attention, despite its importance for practical data analysis. This setting arises from a wide range of applications, for instance, functional magnetic resonance imaging (fMRI) and the financial market contain multivariate time-varying observations. To understand the coupling dynamics of two sets of variables with time-stamped observations or to incorporate temporal structures, a few methods have been proposed. For instance, [13], maximized the auto-correlation of fMRI time series, and [14] proposed to use KCCA to maximize the correlation between the two data sources over a certain time window. Although these approaches do incorporate the temporal dependencies to some extent, the temporal dynamics of consecutive linear transformations (vectors) have not been considered explicitly. Furthermore, these studies focus on performing canonical correlation analysis on two sets of variables with time lags.

A simple method to obtain the dynamic coupling between two sets of variables with time-stamped observation is to apply sparse CCA (for high dimensional data) or KCCA for each timestamp and then compare the vectors. However, in this way, we will lose temporal information and possibly reach the wrong conclusion about temporal dynamics. One such example will be discussed in detail later in this paper. Although one can adopt fused lasso penalty [15] directly, more challenges will be raised in this case. First, CCA for high dimensional data with multiple time-varying observations will become computational expensive (with hundreds of non-convex constraints and many possible local optimums). Temporal incoherence [14] is another severe problem. For example, if we have *w*_1_ and *w*_2_ as our two canonical vectors, according to the definition of CCA problem, the correlation is also maximized by another two vectors *-w*_1_ and *-w*_2_. There is no guarantee that we get the same absolute sign for canonical vectors of two adjacent timestamps even if the data from these two timestamps are same, especially when optimizing non-convex and non-smooth objective functions for these problems.

In order to solve the aforementioned problems, we propose a novel method for inferring the dynamic dependence between two sets of variables. This method integrates the SVD characterized formulation of all solutions of CCA [8] and the fused lasso regularization [15] in a unified optimization framework. We introduce a convex optimization problem which can be solved efficiently. There still exists time incoherence problem in this formulation which we will discuss how to solve later in this paper. We note that since the focus of the paper is on introducing temporal structure in the CCA framework, we will only consider experiments of the first pair of canonical vectors. TDCCA and our algorithm can also be applied in the situations of multiple pairs of canonical vectors.

We summarize our contributions as follows:

- We incorporate temporal dependencies with (sparse) canonical correlation analysis using a convex formulation.
- We propose a fast parallelizable algorithm (Alternating Direction Method of Multipliers) and derive a closed-form ADMM updates to solve the non-smooth objective function.
- Our proposal provides a heuristic method to solve the time incoherence problem that exists in canonical vectors of adjacent timestamps.
- Experimental results on both two different simulated datasets show the effectiveness and accuracy of our method compared with static CCA. The experiments on the real dataset illustrate potential applications of our method for analyzing longitudinal data [16].

## 2. Canonical Correlation Analysis

We first review the standard canonical correlation analysis problem as

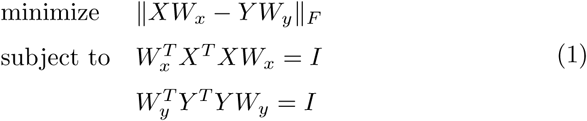

Where 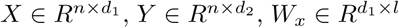 and 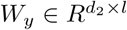. Let *r* = *rank*(*X*), *s* = *rank*(*Y*) and *t* = *min*(*r, s*). *l* is the number of pairs of canonical vectors we attempt to compute. *d*_1_ and *d*_2_ are the dimension of features for *X* and *Y*. *n* is the number of observations. We assume both *X* and *Y* are column centered. Under our temporal setting, *X* and *Y* are the observations at the same time point, see more details later.

Theorem 1 characterizes the solution of (1) by a SVD approach. Let us consider the SVD of X and Y,

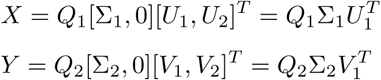

where 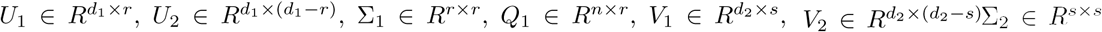 and *Q*_2_ ϵ *R*^*n×s*^. Furthermore, we consider the SVD of 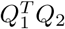 as 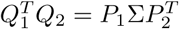 where 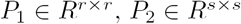 and 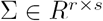. Denote the distinct eigenvalues of 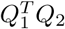 as *σ*_1_ > *σ*_2_ > … > *σ*_*q*_ > 0 with multiplicity for these q eigenvalues being 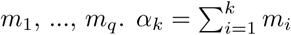

The following theorem from [8] shows the conditions for (*W*_*x*_, *W*_*y*_) which will be used later.

### Theorem 1.

*[8]* If 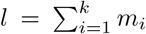 *for some* 1 ≤ *k* ≤ *q, then* (*W*_*x*_, *W*_*y*_) *is a solution of optimization problem* (1) *if and only if*

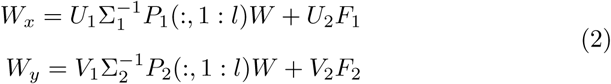

*where W ∈ R*^*l*×*l*^ *is orthogonal*, 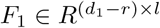 *and* 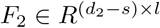 *are arbitrary.*

## 3. Methodology

Before we introduce our proposed method TDCCA, we provide some motivation for our framework, which is related to sparse CCA. If 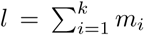 for some 1 ≤ *k* ≤ *q*, the sparse canonical correlation analysis can be stated as solving the following problem,

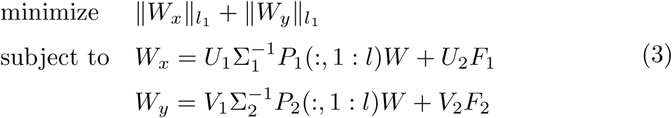

where 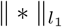 is defined with element-wise *l*_1_ penalty. This formulation is non-convex. An alternative formulation of the above is

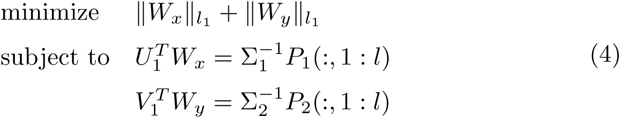

We will further provide a justification for the simplified formulation of sparse CCA problem (4). Denote the optimal value (feasible region) of problem (3) and (4) as *M*_*s*_(Ω_*s*_) and *M* (Ω) respectively.

### Theorem 2.

*If* 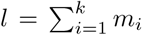 *for some* 1 ≤ *k* ≤ *q, let* 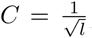 *then CM* ≤ *M*_*s*_ ≤ *M*

*Proof.* First it is easy to see

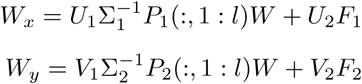

is equivalent to

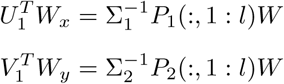

By taking *W* = *I*, we get *M*_*s*_ ≤ *M*.

On the other hand, for *W*_*f*_ and 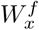 which satisfy (5),

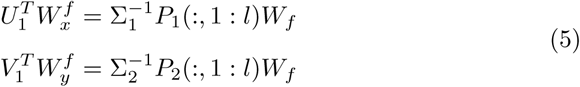

we get

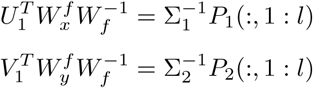

because *W*_*f*_ is orthogonal. This implies 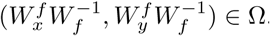. Furthermore, for all *W*_*x*_, *W*_*y*_ ∈ Ω and *W* orthogonal,

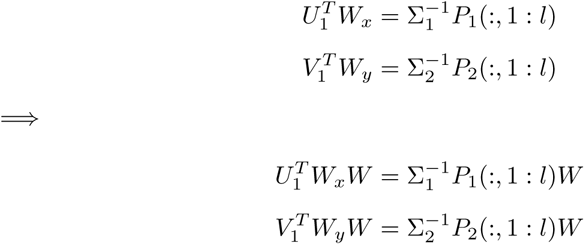

This means (*W*_*x*_*W, W*_*y*_*W, W*) ∈ Ω_*s*_. Specifically, we can take 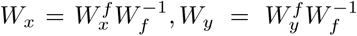 and *W* = *W*_*f*_ for all 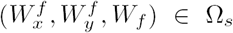. This implies {(*W*_*x*_*W, W*_*y*_*W, W*)|(*W*_*x*_, *W*_*y*_) ∈ Ω, *W* is orthogonal} = Ω_*s*_ and thus problem (3) is equivalent to (6)

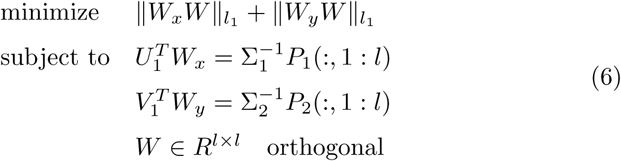

Based on the equivalences of norm of finite dimensional spaces and the orthogonality of *W*, we have

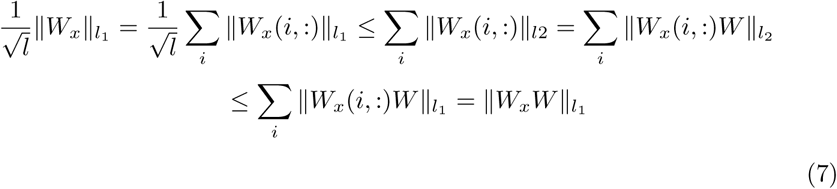

We can get similar result for *W*_*y*_. Inequality (7) implies *CM* ≤ *M*_*s*_ and we already know *M*_*s*_ ≤ *M*, thus *CM* ≤ *M*_*s*_ ≤ *M* where 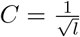

### 3.1 Problem Formulation

Let’s now consider time-dependent views of column centered data 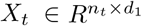 and 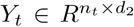 for *t* ∈ [1, 2, …, *T*]. We attempt to analyze dynamic coupling canonical vectors 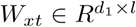 and 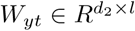 in a CCA framework which incorporate temporal information of these time-stamped observations. It is worth mentioning that our data *X*_*t*_ or *Y*_*t*_ does not have to be the observations from the same time point. They can be selected using a sliding window approach. In particular, a temporal window with length W, is chosen, and within the temporal interval that it spans (from time t=1 to time t=W), the first set of data are selected as *X*_1_ and *Y*_1_. Then, the window is shifted by a step T, and the same data extraction procedure is repeated over the time interval [1 + *T*, *W* +*T*]. This process is iterated until the window spans the end part of the time series. Motivated by Theorem 1 and Theorem 2, we can formulate the problem as

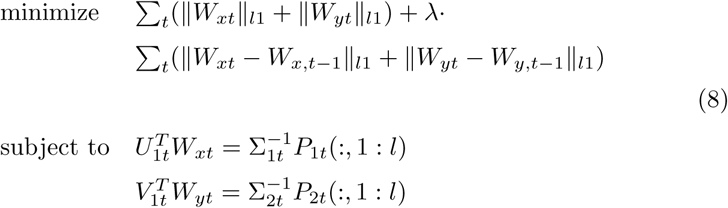

Where

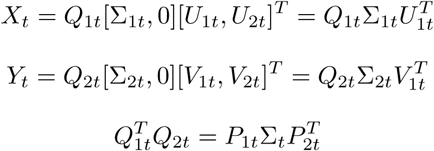

are SVDs for each *t*.

By allowing the relaxation of the constraints, we propose the TDCCA by optimizing the following objective function

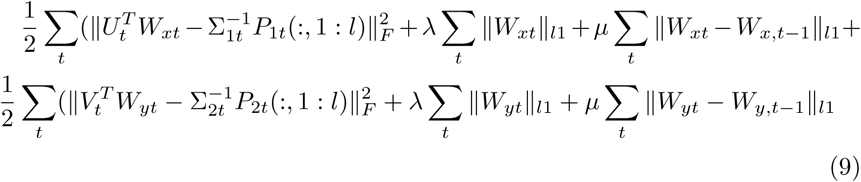

It is clear that the problem we formulate is a convex problem, which avoids the constraints 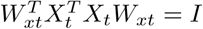 and 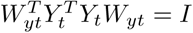 in CCA framework.

### 3.2. Optimization

We present an algorithm to optimize the objective function of TDCCA in (9). The estimation of *W*_*x*_ and *W*_*y*_ can be separated which makes it possible for parallel computing. In addition, the estimation of different pairs of canonical vectors can also be computed in parallel. For the ease of notation, we will ignore the dimension number inside of *P*_1_ which actually represents *P*_1_(:, *ll*) if we try to estimate *ll*-th pair of canonical vectors. Without loss of generality, we will only discuss the algorithm for 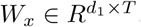, the first pair of canonical vectors.

By separating (9), we get

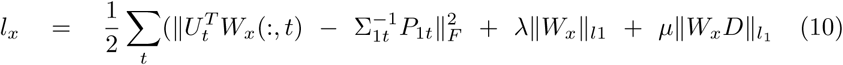

*D* is the time differencing operator, i.e.

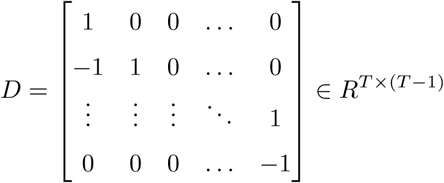

Replacing *W*_*x*_ with 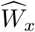 and 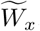, (10) is equivalent to (11)

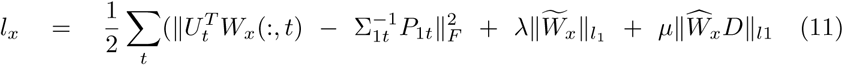

subject to 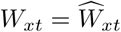 and 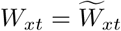

Now we can adopt ADMM [17] to optimize (11). We first write down the augmented Lagrangian for (11),

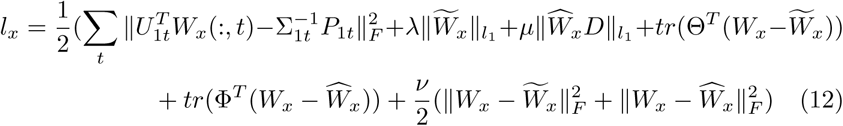

It can be solved by alternatively updating the five variables *W*_*x*_, 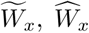, Θ and Φ.

1. Fix 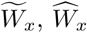, Θ and Φ, we get

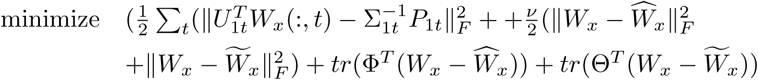

Simply by setting the derivative with respect to *W*_*x*_ to zero, we have

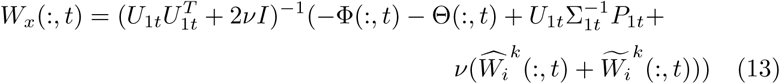
2. Fix *W*_*x*_, 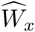, Θ and Φ, the problem is transformed to

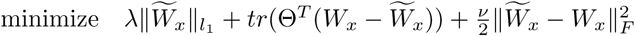

we thus get

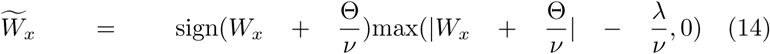
3. Fix *W*_*x*_, 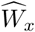, Θ and Φ, (12) becomes

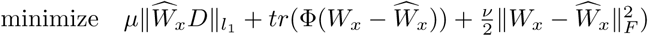

which is equivalent to

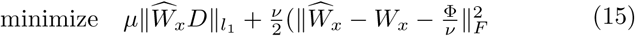

It is a combination of 1-d fused lasso problems which can be solved exactly using dynamic programming method [18] or a taut string principle [19] (both linear time algorithm) in parallel. The similar process can be applied to Θ and Φ.
4. 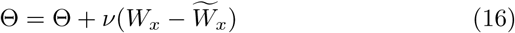
5. 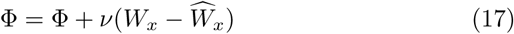

We summarize our algorithm in Algorithm 1 (TDCCA-1). It is worth noting that in our proposed algorithm, step 2 and 3 can run in parallel due to that fact that the computation of 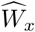 and 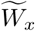 only depends on *W*_*x*_.

### 3.3. Time Incoherence

In (8), there exists a problem called time incoherence. The reason for this problem is that the original constraint in Theorem 1 is 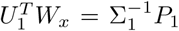(:,1:*l*)*W* where *W* is orthogonal. For *l* = 1, *W* ∈ *R*^1×1^ can be either 1 or −1, which causes the sign ambiguity. In previous section, we ignored the constraint on *W*,

#### Algorithm 1 Algorithm of TDCCA method (TDCCA-1)

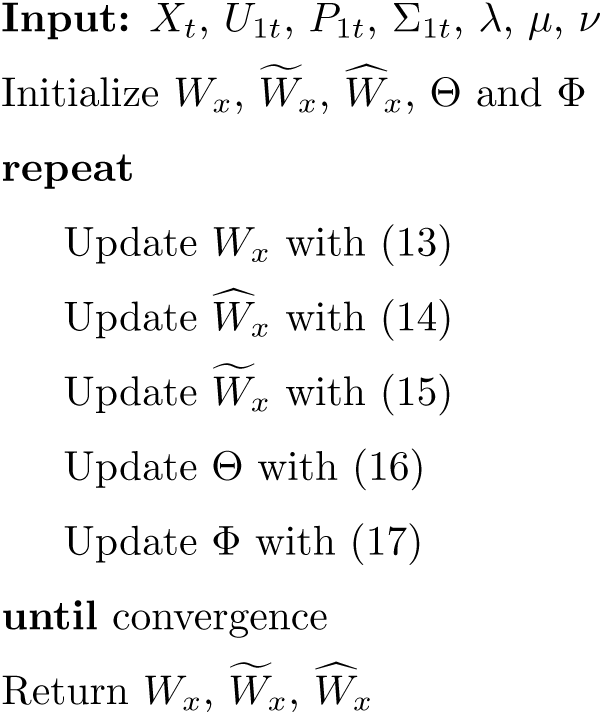

i.e the sign for the case *l* = 1. The problem could be tackled by adding integer variable *b*_*t*_ ∈ {*-*1, 1} and we get a new optimization problem,

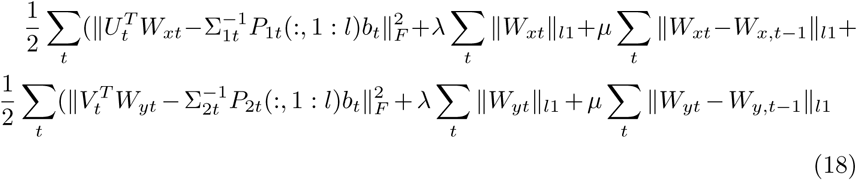

One naive way to solve (18) is to compute the optimal value for every choice of sequence [*b*_1_, …, *b*_*T*_]. The computational burden will increase exponentially. Problem (18) is a non-convex mixed integer problem which is generally hard to solve.

Instead of diving into the non-convex problem, we propose a three-step approach. First, we will use Algorithm 1 with a very small (1e-10 chosen in our experiments) *µ* and thus the temporal difference is not penalized. This step allows us to obtain an initial estimation of *W*_*x*_. Then we will change the sign of *P*_1_ and *P*_2_ according to whether the condition (19) is satisfied. Finally, we run Algorithm 1 again with *µ* chosen by grid search and obtain the final estimations. This method allows us to detect those SVD results with the incoherent sign and thus the temporal consistency is achieved. The intuition behind this approach is that the optimal value of (9) continuously depends on *µ*. The final algorithm is summarized in Algorithm 2.

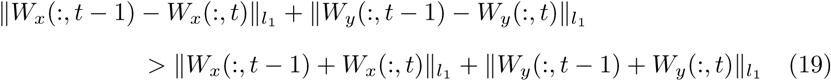

#### Algorithm 2 Algorithm of TDCCA method

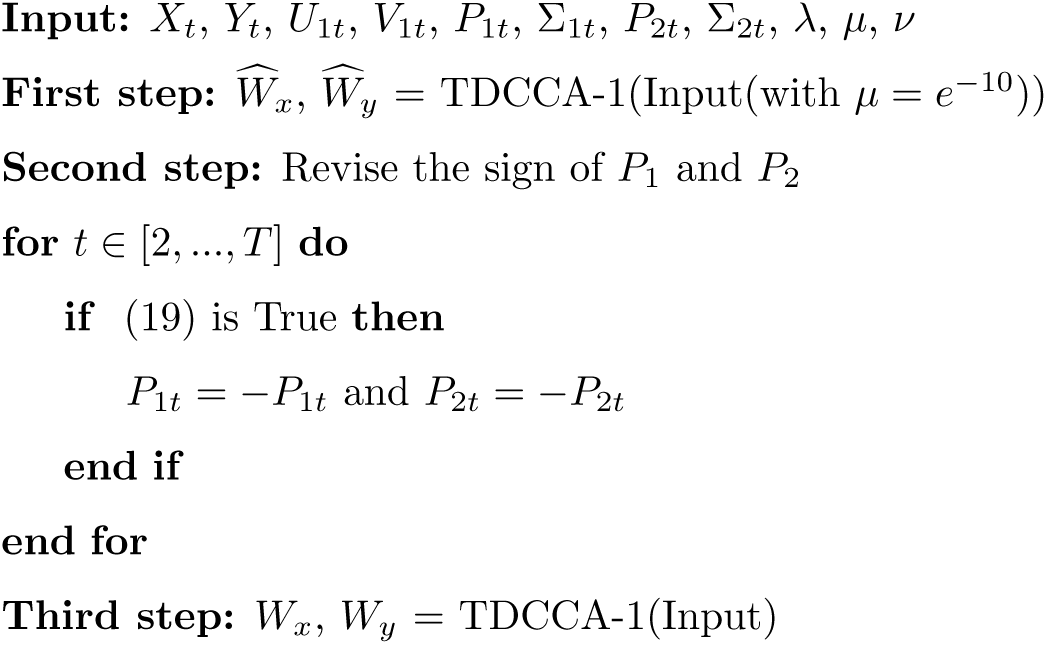

As the Algorithm 1 is computationally efficient, Algorithm 2 is still efficient, considering we will fix *µ* in the first step. Deflation method [6, 20, 7] can also be easily combined with our algorithm after calculating each pair of canonical vectors to acquire multiple pairs of canonical vectors.

### 3.4. Tuning Parameter Selection

In our method, TDCCA contains two tuning parameters *λ* and *µ* which determine the sparsity and continuity (along temporal dimension) of canonical vectors. We propose a cross-validation approach as follows: we partition our data as training and validation data, and then we select the tuning parameters that maximize the canonical correlation on the validation data, plugging the canonical vectors solved from the training data. We also apply the grid search to determine the optimal values of *λ* and *µ*.

### 3.5. Algorithm Analysis

The convergence of ADMM under certain conditions has been analyzed and proven in previous papers [21, 22, 23]. Our optimization problem satisfies the conditions and thus our algorithm is guaranteed to converge to a non-empty solution set if it exists. In each iteration, the major computation burden is in step 1. One iteration of step 2, 3, 4, 5 is *O*(*dT*) where *d* is dimension of feature space. For the first step, the inverse of 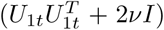 can be precomputed before the iterations. For each iteration, time complexity of step 1 (matrix multiplication) is *O*(*d*^2^*T*). Another feature of our algorithm is both step 1 and 3 can run in parallel along the temporal dimension.

## 4. Experiments

In this section, we evaluate the performance of the proposed method on two simulated datasets and real functional magnetic resonance imaging (fMRI) data. We compare our method with static sparse CCA [6]. For the static sparse CCA, we treat each *t* as an independent problem. We use the R package called PMA for SCCA, which is publicly available at https://cran.r-project.org/web/packages/PMA/index.html. The package for our method is available at https://github.com/xuefeicao/tdcca.

### 4.1. Simulations

We introduce four metrics to measure the accuracy of our estimates in simulations.

- **Correlation Deviation Ratio (CDR)**: This evaluates the capability of our method to recover the true correlation between two sets of variables. It is defined as the ratio of *l*_1_ distance between estimated correlation and true correlation to the true correlation.
- **F1 score**: This measures the ability of our method to capture the true pattern of related variables.
- **Cosine of Angle between estimated and real canonical vectors**: This measures the similarity of our estimation and real canonical vectors. It is defined as the absolute value of cosine angle between two vectors.
- **Temporal Deviation Ratio (TDR)**: The temporal deviation defined as 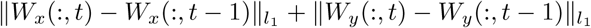, illustrates how much the estimation changes at each time step. This value is the ratio of temporal deviation at change point (in our simulation, for simplicity, only one change point is included) to the average temporal deviation value of all time points. This metric serves the purpose of testing the ability of our method to detect temporal dynamics. We notice that SCCA method does not distinguish the absolute sign of the canonical vectors (*-W* or *W* can both be solutions). To achieve a fair comparison, we alter the canonical vector *W*_*t*_ to *-W*_*t*_ obtained in SCCA method at time *t* if (19) is satisfied. All the results reported in this section are the averages along temporal dimension. We denote *W*_*xt*_ = *W*_*x*_(:, *t*) and 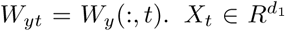 and 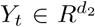. We fix *T* = 100 and *d*_1_ = *d*_2_ for simplicity. Let *d* = *d*_1_ + *d*_2_

#### 4.1.1. Simulation 1

In this simulation, we generate our data according to the following model.

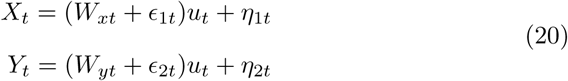

where *u*_*t*_ ∼ *N* (0, 1), *ϵ*_*it*_ ∼ *N* (0, 0.1^2^) and *η*_*it*_ ∼ *N* (0, 0.1^2^) for *i* = 1, 2. For *t* ≤ 100

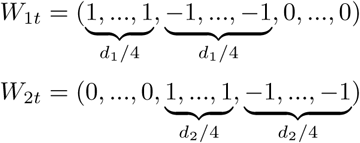

For *t* > 100,

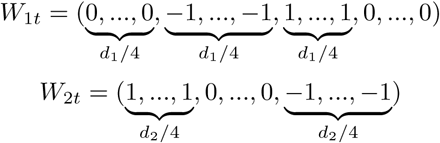

From the model, we can see that for *t* ≤ 50, the first half variables of *X*_*t*_ and second half variables of *Y*_*t*_ are correlated, while for *t* > 50, variables of *X*_*t*_ located in 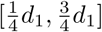 are correlated with variables of *Y*_*t*_ located in 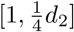 and 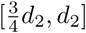 To test our algorithm, we conducted the estimation with different settings:

- *n* = 100, *d* = 40
- *n* = 100, *d* = 100
- *n* = 100, *d* = 400

where *n* is the number of samples. We summarize our results of 50 independent trials in Table 1.

**Table 1:** Simulation 1 results from 50 independent trials

| n | d | T | Method | CDR | F1 | TDR | Cosine of Angle |
| --- | --- | --- | --- | --- | --- | --- | --- |
| 100 | 40 | 100 | TDCCA | 0.0021 | 1.0000 | 97.7767 | 0.9800 |
|  |  |  | SCCA | 0.0025 | 0.9488 | 5.5884 | 0.9693 |
| 100 | 100 | 100 | TDCCA | 0.0008 | 1.0000 | 89.8201 | 0.9730 |
|  |  |  | SCCA | 0.0021 | 0.8192 | 2.1969 | 0.8439 |
| 100 | 400 | 100 | TDCCA | 0.0002 | 0.9977 | 67.5984 | 0.9460 |
|  |  |  | SCCA | 0.0025 | 0.5771 | 1.6001 | 0.6164 |

#### 4.1.2. Simulation 2

In this section, we employ data generated from a more complicated model called single canonical pair model [24]. The model is described in (21). We used the same *W*_*xt*_ and *W*_*yt*_ as in the first simulation.

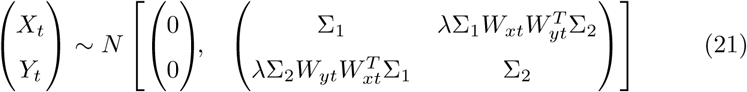

where 0 < *λ* ≤ 1, 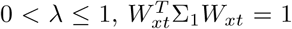 and 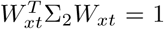. Let *λ* = 0.9. We define Σ_1_ = Σ_2_ = (*σ*_*ij*_)_*ij*_ where *σ*_*ij*_ = *c* × 0.3^|*i-j*|^ which indicates covariance has a certain rate of decay. The scaling factor *c* is obtained by normalization. In addition, we add independent noise 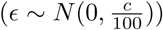 to the generated data. It is easily to verify that for model (21), *W*_*xt*_ and *W*_*yt*_ are first pair of canonical vectors, which maximizes the correlation between 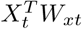 and 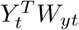 Furthermore, the corresponding correlation is *λ*. We note that our method TDCCA does not depend heavily on the Gaussian covariance assumption. However, in [24], their Sparse CCA method utilizes the model structure (21) 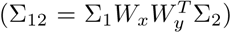 explicitly and then get an estimation of *W*_*x*_ and *W*_*y*_ directly where Σ_12_ is cross-covariance between *X* and *Y*. We used the following settings in this simulation,

- *n* = 100, *d* = 40
- *n* = 100, *d* = 100

where *n* is the number of samples. Table 2 showed the averaged results of 50 independent trials.

**Table 2:** Simulation 2 results from 50 independent trials

| n | d | T | Method | CDR | F1 | TDR | Cosine of Angle |
| --- | --- | --- | --- | --- | --- | --- | --- |
| 100 | 40 | 100 | TDCCA | 0.0599 | 0.9896 | 74.8900 | 0.9297 |
|  |  |  | SCCA | 0.0398 | 0.8177 | 2.2071 | 0.9039 |
| 100 | 100 | 100 | TDCCA | 0.0339 | 0.9647 | 53.1478 | 0.9642 |
|  |  |  | SCCA | 0.0412 | 0.7317 | 1.3389 | 0.7692 |

#### 4.1.3. Simulation Results

We show results for both models in Table 1 and Table 2. In terms of F1 score, Temporal Deviation Ratio and Cosine of Angle between estimated and real canonical vectors, our TDCCA approach significantly outperforms SCCA method (i.e the CCA method without considering time series structure). The F1 score of TDCCA stays above 0.9 in different settings of two simulations while the F1 score of SCCA can be less than 0.6 when the ratio 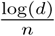 increases. The cosine of Angle of our approach is close to 1 which indicates the high similarity between estimated and real canonical vectors. In addition, the temporal deviation ratio of TDCCA is up to 40 times higher than the SCCA method, which shows a big advantage of TDCCA in detecting change points. Furthermore, our estimated value of correlation is closer to the true correlation of simulated data than the SCCA method.

### 4.2. Canonical Correlation Variation for resting state fMRI

In this section, we apply our method on the resting state fMRI data and we propose that the canonical correlation variation (CCV), a new metric obtained from our method can provide clues for the connectivity patterns that transfer and present aging features. Canonical correlation variability (CCV) is defined as the standard deviation of time-dependent canonical correlation from our method.

Recent work has shown that functional connectivity is temporally dynamic and functional connectivity fluctuates across shorter time-windows for resting state fMRI [25, 26]. Unlike conventional FC analysis, which assumes static connectivity over several minutes, the dynamic functional connectivity variation (FCV) is calculated as the standard variation of the dynamic FC series. In this approach, the stability of the FC fluctuation over time is quantitatively measured and compared between brain region pairs. The basic sliding window framework has been used widely and is repeatedly applied by researchers to investigate how functional brain dynamics relates to our cognitive abilities [27]. Age-related dynamic pattern of functional connectivity has been also explored in [28, 22, 29].

Canonical correlation is another way to characterize the strength of the functional connectivity for each region pair which has been used to construct a region-level functional connectivity network for predicting major depressive disorder [30]. In our experiment, two groups of individuals (N = 156, ages 22–25 for the first group; N = 226, ages 31–35 for the second group) were recruited from the public data of the Human Connectome Project (http://www.humanconnectomeproject.org/). In particular, we are interested in the connectivity patterns inside of default mode network (DMN) which contains Precuneus (pC), Posterior cingulate (PCC), Ventral anterior cingulate (vACC), and Medial prefrontal cortex (mPFC). For each subject, we used a fixed-length rectangle window (width = 60 TRs) and the window was shifted by 2 TRs. These parameters are chosen based on the rule of choosing parameters for dynamic functional connectivity [27]. Thus we can obtain the time-dependent canonical correlation estimated for each pair of ROIs from our method for every subject from two groups.

Additionally, we use two popular measurements of connectivity pattern: static functional connectivity (FC), functional connectivity variation (FCV). As a baseline method, we compute the sparse canonical correlation for each rectangle window and calculate its standard deviation which we will call it CCVB in the remaining paper. The canonical correlation coefficient (CCC) for the entire time series is also included for each subject. These features are summarized in table 3.

**Table 3:** Features of connectivity pattern used in our experiment

| Method | Description |
| --- | --- |
| CCV | Canonical Correlation Variation calculated from TD-CCA |
| FC | Static Functional Connectivity (Fisher-transformed correlations) of entire time series |
| FCV | Functional Connectivity Variation using sliding window approach |
| CCVB | Canonical Correlation Variation calculated from SCCA for each sliding window |
| CCC | Canonical Correlation Coefficient (Fisher-transformed correlations) of entire time series |

Table 4 shows the p–value of two sample t-test of different features for each pair of ROIs compared between two different groups. Multiple testing correction is performed using the FDR method [31]. From table 4, we can see the only one significant difference for metrics calculated based on two different groups are from our proposed canonical correlation variation measurement. It is between vACC and PCC. Figure 1 illustrates the group differences of different features for the ROI pair with a significant p–value. The values are scaled for better visualization. It shows CCV (between vACC and PCC) in the age group 31– 35 is higher than the age group 22–25. This example shows promising results by applying CCV as a novel way to measure dynamic functional connectivity pattern using resting state fMRI.

**Table 4:** Adjusted p–value for each pair of ROIs chosen from default mode network. Values in bold represent significant p–value with threshold 0.05.

| ROI 1 | ROI 2 | CCV | FC | FCV | CCVB | CCC |
| --- | --- | --- | --- | --- | --- | --- |
| vACC | pC | 0.5197 | 0.8092 | 0.9452 | 0.9498 | 0.7419 |
| vACC | PCC | 0.0138 | 0.9725 | 0.9452 | 0.4273 | 0.7419 |
| PCC | pC | 0.5197 | 0.8092 | 0.9452 | 0.2464 | 0.7419 |
| mPFC | vACC | 0.4224 | 0.9725 | 0.9452 | 0.4273 | 0.7419 |
| mPFC | pC | 0.1593 | 0.8092 | 0.9452 | 0.2464 | 0.7419 |
| mPFC | PCC | 0.5197 | 0.9725 | 0.9452 | 0.2464 | 0.7419 |

**Figure 1:**
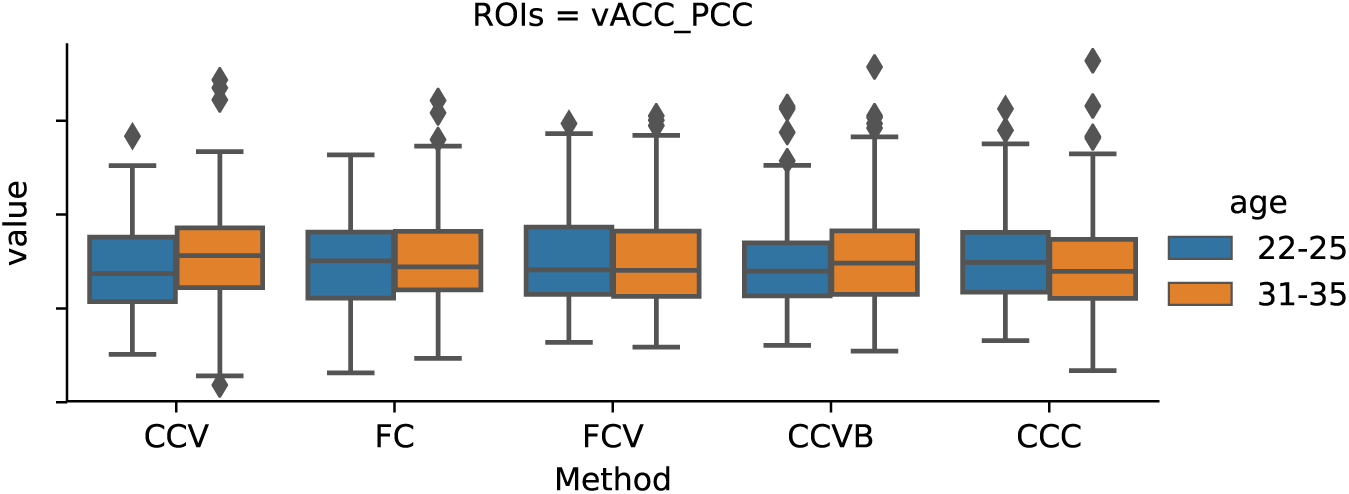

### 4.3. Task-based fMRI: motor task detection and feature extraction

In this section, we apply the TDCCA method to analyze task-related fMRI motor data obtained from the Human Connectome Project [32]. This example will show how our method can spot change points due to different tasks.

This motor task fMRI is composed of five most basic motor tasks including tapping left/right fingers, squeezing left/right toes and moving tongue. Participants were presented with visual cues which asked them to either tap their left or right fingers, or squeeze their left or right toes, or move their tongue to map motor areas. Each block of a movement type lasted 12 seconds (5 movements), and was preceded by a 3-second cue.

Based on the prior scientific findings on the motor task experiment [33], we select six brain regions corresponding to these different tasks according to MNI coordinates: left/right hand coordinates (±41, 20, 62), left/right foot coordinates(±6, 26, 76), tongue coordinates (±55, 4, 26), thalamus (MNI: −12, −13, 7). We extracted voxels around these coordinates, depending on the availability of voxels centered around these coordinates. Thus we combine data from these six regions as our one set of data. We set the length of the sliding window as 20 TRs according to the length of each task and the window was shifted by 1 TR (0.72 s). Our TDCCA method is applied to a pair of subjects. We should mention that brain electrical activity is not directly measured, instead, the human hemodynamic responses to the short period of neural activity are delayed in time. Thus fMRI measures the subsequent demand for oxygenated blood that follows about several seconds after the neuronal activations [34].

We estimate the leading canonical vectors and elaborate on how the *W*_*x*_ varies with time periods of the different task activation. We also compared the SCCA (sparse CCA method) with TDCCA. Figure 2 shows the scaled temporal deviation 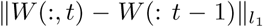 estimated from two methods. Figure 2 elaborates the ability of our approach in detecting temporal dynamics. From the results of SCCA and original BOLD signal, one can barely see the dynamic change point for different motor tasks. However, our method detects six significant shift of tasks and a clear time delay from task commands to the peak of temporal deviation.

**Figure 2:**
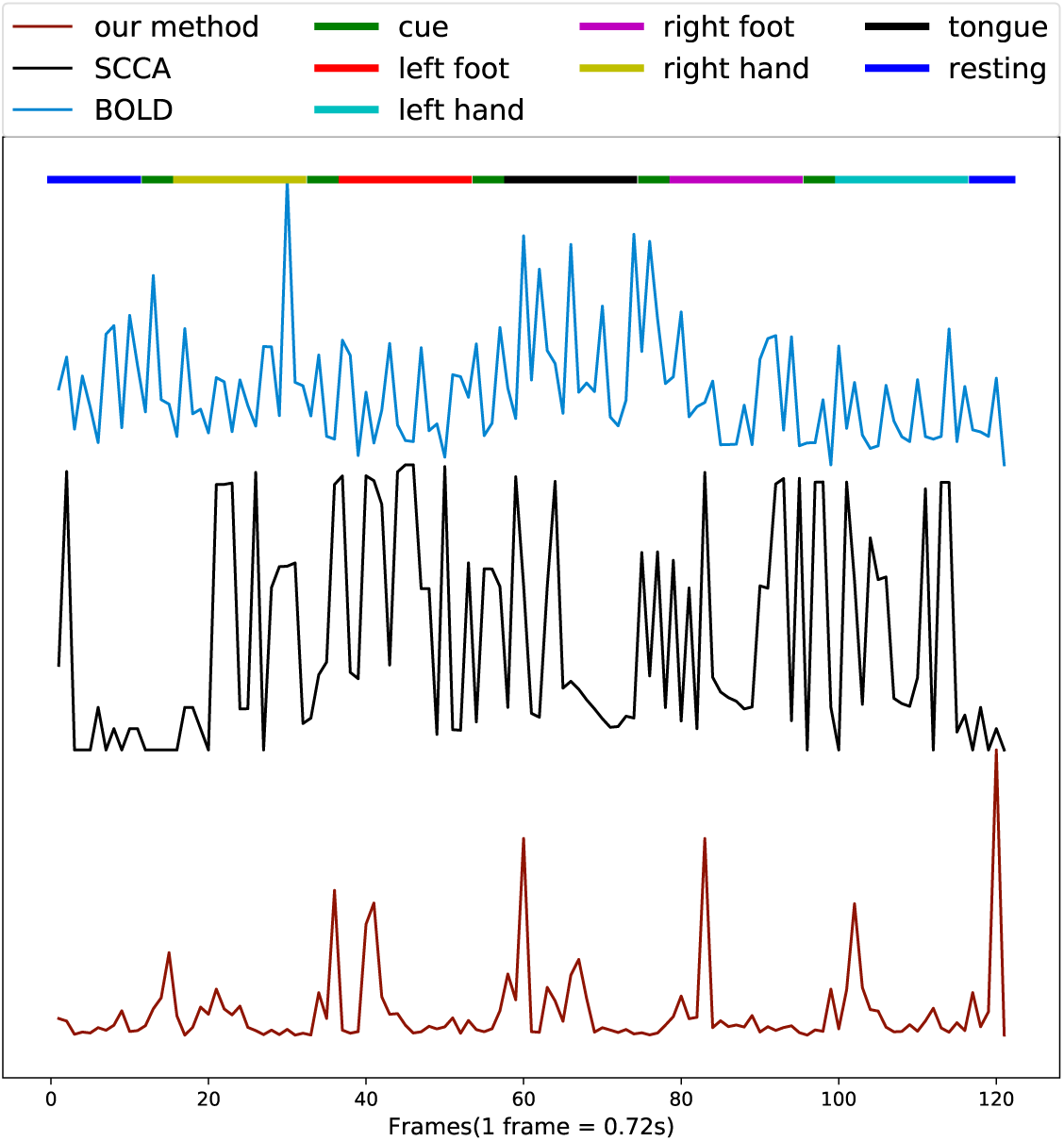
Plot of the temporal deviation of TDCCA, SCCA and original BOLD signal, in which TDCCA detect six significant shift of tasks. The straight line above the plot uses different colors to indicate the different tasks during the experiment.

## 5. Conclusions and future work

In this paper, a convex framework for combining temporal structure with canonical correlation analysis is proposed. The proposed framework incorporates temporal information explicitly. Furthermore, our algorithm is computationally efficient with guaranteed convergence and has the advantage of parallel computing. Finally, we introduce a heuristic method to solve the time incoherence problem without using a mixed integer optimization algorithm. The proposed method outperforms the (static) sparse CCA algorithm both in accuracy and ability to recover temporal variations. Our proposed canonical correlation variation (CCV) can also provide clues for brain connectivity patterns. Our method introduces an additional tool to determine change points and extract critical features in multivariate analysis. In future work, we will explore the theoretical property of our proposed algorithm. It would be also promising to apply our method to analyze multivariate longitudinal data from medical images.

## References

[1] H. Hotelling, Relations between two sets of variates, Biometrika 28 (3/4) (1936) 321–377.

[2] S. Akaho, A kernel method for canonical correlation analysis, in: International Meeting on Psychometric Society, 2001, 2001.

[3] T. Melzer, M. Reiter, H. Bischof, Nonlinear feature extraction using generalized canonical correlation analysis, in: International Conference on Artificial Neural Networks, Springer, 2001, pp. 353–360.

[4] S. Waaijenborg, P. C. V. de Witt Hamer, A. H. Zwinderman, Quantifying the association between gene expressions and dna-markers by penalized canonical correlation analysis, Statistical applications in genetics and molecular biology 7 (1).

[5] D. M. Witten, R. Tibshirani, T. Hastie, A penalized matrix decomposition, with applications to sparse principal components and canonical correlation analysis, Biostatistics 10 (3) (2009) 515–534.

[6] D. M. Witten, R. J. Tibshirani, Extensions of sparse canonical correlation analysis with applications to genomic data, Statistical applications in genetics and molecular biology 8 (1) (2009) 1–27.

[7] D. R. Hardoon, J. Shawe-Taylor, Sparse canonical correlation analysis, Machine Learning 83 (3) (2011) 331–353.

[8] D. Chu, L.-Z. Liao, M. K. Ng, X. Zhang, Sparse canonical correlation analysis: new formulation and algorithm, IEEE transactions on pattern analysis and machine intelligence 35 (12) (2013) 3050–3065.

[9] O. Friman, J. Cedefamn, P. Lundberg, M. Borga, H. Knutsson, Detection of neural activity in functional mri using canonical correlation analysis, Magnetic Resonance in Medicine 45 (2) (2001) 323–330.

[10] D. R. Hardoon, S. Szedmak, J. Shawe-Taylor, Canonical correlation analysis: An overview with application to learning methods, Neural computation 16 (12) (2004) 2639–2664.

[11] J. Rupnik, J. Shawe-Taylor, Multi-view canonical correlation analysis, in: Conference on Data Mining and Data Warehouses (SiKDD 2010), 2010, pp. 1–4.

[12] T.-K. Kim, S.-F. Wong, R. Cipolla, Tensor canonical correlation analysis for action classification, in: Computer Vision and Pattern Recognition, 2007. CVPR’07. IEEE Conference on, IEEE, 2007, pp. 1–8.

[13] O. Friman, M. Borga, P. Lundberg, H. Knutsson, Exploratory fmri analysis by autocorrelation maximization, NeuroImage 16 (2) (2002) 454–464.

[14] F. Bießmann, F. C. Meinecke, A. Gretton, A. Rauch, G. Rainer, N. K. Logothetis, K.-R. Müller, Temporal kernel cca and its application in multimodal neuronal data analysis, Machine Learning 79 (1-2) (2010) 5–27.

[15] R. Tibshirani, M. Saunders, S. Rosset, J. Zhu, K. Knight, Sparsity and smoothness via the fused lasso, Journal of the Royal Statistical Society: Series B (Statistical Methodology) 67 (1) (2005) 91–108.

[16] G. Verbeke, S. Fieuws, G. Molenberghs, M. Davidian, The analysis of multivariate longitudinal data: A review, Statistical methods in medical research 23 (1) (2014) 42–59.

[17] S. Boyd, N. Parikh, E. Chu, B. Peleato, J. Eckstein, et al., Distributed optimization and statistical learning via the alternating direction method of multipliers, Foundations and Trends*Q*R 1–122. in Machine learning 3 (1) (2011)

[18] N. A. Johnson, A dynamic programming algorithm for the fused lasso and l 0-segmentation, Journal of Computational and Graphical Statistics 22 (2) (2013) 246–260.

[19] P. L. Davies, A. Kovac, Local extremes, runs, strings and multiresolution, Annals of Statistics (2001) 1–48.

[20] J. Shawe-Taylor, N. Cristianini, Kernel methods for pattern analysis, Cambridge university press, 2004.

[21] B. He, X. Yuan, On the o(1/n) convergence rate of the douglas–rachford alternating direction method, SIAM Journal on Numerical Analysis 50 (2) (2012) 700–709.

[22] C. Chen, B. He, Y. Ye, X. Yuan, The direct extension of admm for multi-block convex minimization problems is not necessarily convergent, Mathematical Programming 155 (1-2) (2016) 57–79.

[23] M. Hong, Z.-Q. Luo, On the linear convergence of the alternating direction method of multipliers, Mathematical Programming 162 (1-2) (2017) 165–199.

[24] M. Chen, C. Gao, Z. Ren, H. H. Zhou, Sparse cca via precision adjusted iterative thresholding, arXiv preprint arXiv:1311.6186.

[25] C. Chang, G. H. Glover, Time–frequency dynamics of resting-state brain connectivity measured with fmri, Neuroimage 50 (1) (2010) 81–98.

[26] D. A. Handwerker, V. Roopchansingh, J. Gonzalez-Castillo, P. A. Bandettini, Periodic changes in fmri connectivity, Neuroimage 63 (3) (2012) 1712–1719.

[27] M. G. Preti, T. A. Bolton, D. Van De Ville, The dynamic functional connectome: state-of-the-art and perspectives, Neuroimage 160 (2017) 41–54.

[28] T. M. Madhyastha, T. J. Grabowski, Age-related differences in the dynamic architecture of intrinsic networks, Brain connectivity 4 (4) (2014) 231–241.

[29] Y. Chen, Y.-n. Liu, P. Zhou, X. Zhang, Q. Wu, X. Zhao, D. Ming, The transitions between dynamic micro-states reveal age-related functional network reorganization, Frontiers in Physiology 9.

[30] J. Kang, F. D. Bowman, H. Mayberg, H. Liu, A depression network of functionally connected regions discovered via multi-attribute canonical correlation graphs, NeuroImage 141 (2016) 431–441.

[31] Y. Benjamini, Y. Hochberg, Controlling the false discovery rate: a practical and powerful approach to multiple testing, Journal of the Royal statistical society: series B (Methodological) 57 (1) (1995) 289–300.

[32] D. C. Van Essen, S. M. Smith, D. M. Barch, T. E. Behrens, E. Yacoub, K. Ugurbil, W.-M. H. Consortium, et al., The wu-minn human connectome project: an overview, Neuroimage 80 (2013) 62–79.

[33] B. Thomas Yeo, F. M. Krienen, J. Sepulcre, M. R. Sabuncu, D. Lashkari, M. Hollinshead, J. L. Roffman, J. W. Smoller, L. Zöllei, J. R. Polimeni, et al., The organization of the human cerebral cortex estimated by intrinsic functional connectivity, Journal of neurophysiology 106 (3) (2011) 1125–1165.

[34] C. H. Liao, K. J. Worsley, J.-B. Poline, J. A. Aston, G. H. Duncan, A. C. Evans, Estimating the delay of the fmri response, NeuroImage 16 (3) (2002) 593–606.

